# Source-independent enrichment of light lanthanides: microbial mobilization, selective uptake, and intracellular storage

**DOI:** 10.64898/2026.03.03.709224

**Authors:** Linda Gorniak, Sophie M. Gutenthaler-Tietze, Alina Lobe, Lena J. Daumann, Robin Steudtner, Thorsten Schäfer, Frank Steiniger, Martin Westermann, Kirsten Küsel, Carl-Eric Wegner

## Abstract

Poorly soluble lanthanide minerals pose challenges for both a sustainable extraction of lanthanides as key resources for decarbonization and lanthanide-dependent microbial metabolism. Microbial use of lanthanides is widespread, yet bacteria’s preference for light lanthanides requires differentiation mechanisms that enable downstream utilization. Whether lanthanide discrimination occurs during access, mobilization, uptake, or intracellular processing is mostly unknown and likely controlled by habitat and bioavailability. We studied microbial lanthanide mobilization and uptake from different lanthanide minerals, an alloy, and pure lanthanide compounds. *Beijerinckiaceae* bacterium RH AL1 served as a model organism for an integrated approach combining transcriptomics, analytics, and electron microscopy. This facultative methylotroph depends on light lanthanides for methanol oxidation and forms periplasmic lanthanide deposits. AL1 grew with all tested lanthanide sources and selectively enriched light lanthanides independent of source type, overall lanthanide content, and the proportion of light lanthanides. Transcriptomics revealed that the type of lanthanide source significantly influenced gene expression beyond lanthanide utilization. Lanthanide discrimination in *Beijerinckiaceae* bacterium RH AL1 is a multilayered process rooted in the complementary action of chelation, uptake mechanisms, and periplasmic storage. Adaptations that increase lanthanide bioavailability transform mineral-bound lanthanides into shared resources within microbial communities, with implications for sustainable lanthanide use.

## INTRODUCTION

The interest in microbial lanthanide (Ln, lanthanides [Lns]) utilization intensifies, as Lns (the f-elements, including lanthanum [La-Lu], **Table S1**) are key resources for the ongoing green energy transition [1] and classified as critical resources by government agencies [2]. Abundant in nature, Lns are rarely enriched and co-occur, requiring complex and cost-intensive separation and refinement processes. Lns play an important role in microbial carbon cycling, especially in C_1_-metabolism, and likely have broader roles in microbial metabolism [3–6]. We are only beginning to unravel how microbes mobilize, take up, and distinguish between Lns, all of which can facilitate sustainable Ln use.

Known biomolecules directly involved in Ln sensing, utilization, and transport (summarized as the lanthanome [7]) include the Ln-binding proteins lanmodulin (LanM) [8], lanpepsy [9], landiscernin (LanD) [10], as well as the Ln-utilization and transport (*lut*-) [8, 11, 12] and methylolanthanin (a Ln-binding metallophore) uptake (*mlu*-/*mll*-)clusters [13]. Lanmodulin attracted significant attention, enabling the development of a high-sensitivity Ln sensor and a pilot-scale platform for selective Ln recycling [7, 14]. Known Ln-dependent enzymes rely on pyrroloquinoline quinone as cofactor and function as alcohol dehydrogenases (ADH). Ln-dependent Xox-type methanol dehydrogenases (MDH) represent key enzymes for methanol oxidation in gram-negative bacteria. Transcript abundances of *xoxF* significantly exceed those encoding Ca-dependent Mxa-type MDHs in the surface ocean [15].

Lns co-occur, are chemically similar, and commonly divided into light (La-Eu) and heavy (Gd-Yb) Lns [16, 17]. Only light Lns facilitate Ln-dependent growth [18]. Although heavy Lns have been observed to be taken up and stored [5], no function in microbial metabolism has yet been assigned to them. It is currently unclear if discrimination between light and heavy Lns occurs during mobilization, uptake, or intracellular processing. Studies investigating Ln-dependent metabolism have mainly used soluble Ln salts, often at higher concentrations, which only distantly mimic natural conditions. Lns are typically present as poorly soluble minerals, including monazite and xenotime (phosphate minerals), as well as bastnaesite (a carbonate mineral) [19–22], which are primary Ln sources for mining [23]. Monazite has been the focus of various studies on Ln recovery via bioleaching approaches using known phosphate-solubilizing bacteria (PSB) and fungi [24].

Ln bioavailability depends on the environment. Consequently, mechanisms for Ln mobilization, discrimination, and uptake/storage likely differ between taxa depending on their habitat. In this study, we chose *Beijerinckiaceae* bacterium RH AL1 (strain AL1) to investigate the mobilization of Lns from (i) Ln-containing minerals and the alloy ferrocerium, as well as (ii) poorly soluble Ln forms (La_2_O_3_, Nd_2_O_3_, LaPO_4_). Different mineral types with different Ln contents and proportions of light and heavy Ln elements were tested, including apatite, monazite, and xenotime (phosphate minerals), bastnaesite (a carbonate), loparite (an oxide), and gadolinite (a silicate). The tested minerals differed by recalcitrance, defining different levels of Ln accessibility. Strain AL1 was previously isolated from early-industrial soft-coal slags [25, 26]. It is a facultative methylotroph that naturally depends on light Lns for methanol oxidation. It contains only a single *xoxF* gene in its genome and no *mxaF* homologue. The strain is of particular interest for biotechnological applications, as it has previously been shown to take up Lns and form periplasmic Ln accumulations located at the cell poles, near polyhydroxyalkanoate (PHA) granules [5, 27]. We were interested in whether Ln mobilization, uptake, and intracellular storage are selective processes, and at which stage discrimination between different Lns occurs. Making Lns more bioavailable is a requirement for Lns to play a broader role in microbial communities and microbial physiology.

## MATERIALS AND METHODS

### Sourcing Ln-containing minerals and ferrocerium

Used minerals and ferrocerium have been obtained through colleagues (bastnaesite [origin: Nam Xe, Vietnam; 1983], loparite [USSR, 1988], gadolinite [Norway, 1958], apatite [Finland, Kola Peninsula, Institute of Chemistry and Technology of Rare Elements and Mineral Resources (IChTREMS); 1985]) and retailers (xenotime and monazite [Etsy, TheGlobalStone]. The ferrocerium was donated [Treibacher Industrie AG, Althofen, Austria]).

### Analytical investigation of Ln sources

Elemental analysis (C, H, N, S) was performed at the analytical facility of the Department of Chemistry and Pharmacy, Ludwig Maximilian University of Munich (LMU), Germany, using a Vario EL cube instrument (Elementar Analysensysteme GmbH, Langenselbold, Germany). Inductively coupled plasma mass spectrometry (ICP-MS) was performed on milled and acid-digested minerals at the analytical facility of the Helmholtz Center Dresden-Rossendorf, Germany, using an iCAP RQ ICP-MS device (Thermo Fisher Scientific, Frankfurt, Germany). More information is provided in the **supplementary material**. Mineral compositions were verified by scanning electron microscopy (SEM) coupled to energy-dispersive X-ray spectroscopy (EDX) with a Helios Nanolab G3 Dual Beam UC (FEI, part of ThermoFisher Scientific, Darmstadt, Germany) coupled to an X-Max 80 SDD detector (Oxford Instruments GmbH, Wiesbaden, Germany). The mineral samples were placed on a brass sample carrier with an adhesive carbon foil. A ME-020 high-vacuum coating system (Leica Microsystems [formerly BAL-TEC], Wetzlar, Germany) was used to coat the samples with a conducting carbon film. Measurements were conducted at the Analytic Facility of the Department of Chemistry and Pharmacy at LMU.

### Cultivation

Cultivation with *Beijerinckiaceae* bacterium RH AL1 was done as outlined before [5] in MM2 medium [28] with methanol as the carbon source (123 mM, 0.5% [v/v]) and 0.1% [v/v] of trace element solution no. 1 [29]. Two sets of incubations were done with each condition (= different Ln source) in triplicates. For the first set, La or Nd (1 µM) were supplied in different forms: as (i) chlorides, (ii) oxides, or (iii) phosphate (La only). For the second, Ln-containing minerals (apatite, bastnaesite, gadolinite, loparite, monazite, or xenotime; 5 mg [milled] mineral per culture [140 mL], 0.26 - 16.14 µmol Ln) or the alloy ferrocerium (one piece of 145.27 ± 2.48 mg per culture [140 mL], 770.55 µmol Ln) served as Ln sources. RNA extraction samples were collected during late exponential growth. Cells were harvested and stored at −80°C until further processing. Controls with methanol, but without added Lns, were run in parallel to detect Ln leaching from glassware and carbon source carryover from pre-cultures. Additional controls with Ln minerals or ferrocerium, but no methanol, were used to verify that AL1 did not utilize carbon present in the Ln sources. More information is provided in the **supplementary material**.

### Supernatant analysis

Supernatant samples were collected from biotic samples during biomass harvest for downstream RNAseq analysis. We carried out abiotic incubations in MM2 medium adjusted to pH 5 or 3 to mimic the pH decrease observed during biotic incubations. We matched sampling time points from those to the biotic incubations. Filtered (0.22 µm PES) supernatant samples were analyzed with an 8900 Triple Quadrupole ICP-MS device (Agilent Technologies, Waldbronn, Germany) coupled to an autosampler prepFAST4DX System (Elemental Scientific, Mainz, Germany).

### Electron microscopic investigations of biomass samples

Biomass samples from replicated incubations and matching timepoints (**Figure S3**) were collected via centrifugation (2 ✕ [10 min., 10,000 ✕ g at room temperature]). Sample preparation and subsequent analysis through transmission electron microscopy (TEM) and EDX analyses were done as described previously [5, 27]. EDX spectra have been deconvoluted using the Quantax software (Bruker, Berlin, Germany).

### RNA extraction, mRNA enrichment, library preparation, and sequencing

Late-exponential biomass samples were subjected to total RNA extraction, mRNA enrichment, and RNAseq library preparation as previously described [5, 27]. Sequencing libraries were equimolarly pooled. Illumina sequencing (2 ✕ 100 bp, paired-end) with a NovaSeq 6000 instrument and an SP flowcell (Illumina, San Diego, California, USA) was carried out by the sequencing core facility of the Leibniz Institute on Aging - Fritz Lipmann Institute (Jena, Germany).

### Sequence data pre-processing and differential gene expression analysis

The quality of raw and trimmed sequences was checked with *fastQC* (v0.11.9) [30]. Data pre-processing has been outlined before [5, 6] and is covered in the **supplementary material**. Differential gene expression analysis (DGEA) was performed with the R software environment (v.4.2.1) [31] and the package *edgeR* (v.3.42.4) [32] together with *limma* (v3.50.0) [33], *mixOmics* (v6.18.1) [34], *HTSFilter* (v1.34.0) [35], and *bigPint* (v1.10.0) [36]. Genes featuring changes in gene expression above |0.58| log_2_FC (fold change), gene expression values higher than 4 (log_2_CPM, counts per million), and FDR (false discovery rate)-adjusted *p*-values below 0.05 were considered for further analysis.

### Weighted Gene Correlation Network Analysis (WGCNA)

Pre-processed RNAseq data were filtered based on expression (log_2_CPM values > 4) and normalized with *edgeR*. Co-expression networks were generated using the package *WGCNA* (v.1.73) [37, 38], including relevant dependencies. A soft threshold power of 15 was chosen for network construction based on the scale-free fit index and mean connectivity at various soft-thresholding powers. Genes with similar expression patterns were grouped into nine modules, and correlations between module eigengenes and selected physiological traits were calculated.

### Figure generation

Figures have been generated using the R software framework (v4.2.1), taking advantage of the following packages: *ggplot2* (v3.3.6) [39], *gplots* (v3.1.3) [40], *ggpubr* (v0.4.0) [41], *cowplot* (v1.1.1) [42], and *upsetR* (v1.4.0) [43], including their respective dependencies. Figures have been finalized with *inkscape* (https://inkscape.org/).

### Data availability

The generated RNAseq data sets can be accessed through EBI/ENA ArrayExpress (https://www.ebi.ac.uk/biostudies/arrayexpress/studies/E-MTAB-15640). Further details regarding data processing are available through GitHub (https://github.com/wegnerce/gorniak_et_al_2026). A reproducible, snakemake-based [44, 45] workflow for RNAseq data processing can be retrieved from GitHub (https://github.com/wegnerce/smk_rnaseq, release v0.1).

## RESULTS

### Characterization of different Ln minerals

We examined different types of Ln-containing minerals, including phosphates (apatite, monazite, xenotime), an oxide (loparite), a silicate (gadolinite), and a carbonate (bastnaesite), through EDX and ICP-MS, before using them as Ln sources in incubation experiments (Figure 1A). The determined elemental compositions matched the respective sum formulas, with varying degrees of impurity (Figure 1B**, Table S3**). Monazite featured the highest Ln content (Figure 1A**+B**) among the selected minerals (457.39 µg/mg Ln). Lower Ln concentrations were detected in loparite, xenotime, and bastnaesite, ranging from 132.64 µg/mg (bastnaesite) to 168.10 µg/mg (loparite). Only 38.07 µg/mg and 7.429 µg/mg Lns were detected in gadolinite and apatite. We also considered the alloy ferrocerium as a Ln source, which was primarily composed of cerium (47.1% ± 0.2%), lanthanum (26.7% ± 0.8%), and iron (21.3% ± 0.9%), as determined by ICP-MS (**Table S4**). *Beijerinckiaceae* bacterium RH AL1 was previously shown to require light Lns (La - Nd) for methylotrophic growth. La - Nd made up the majority of Lns (92.5% - 99.3%) in most minerals (monazite, bastnaesite, loparite, and apatite) and ferrocerium, with Ce (44.9% - 62.0% of total Ln) and La (23.5% - 35.2% of total Ln) being especially abundant. In gadolinite and xenotime samples, higher proportions of heavy Lns were detected.

**Figure 1.**
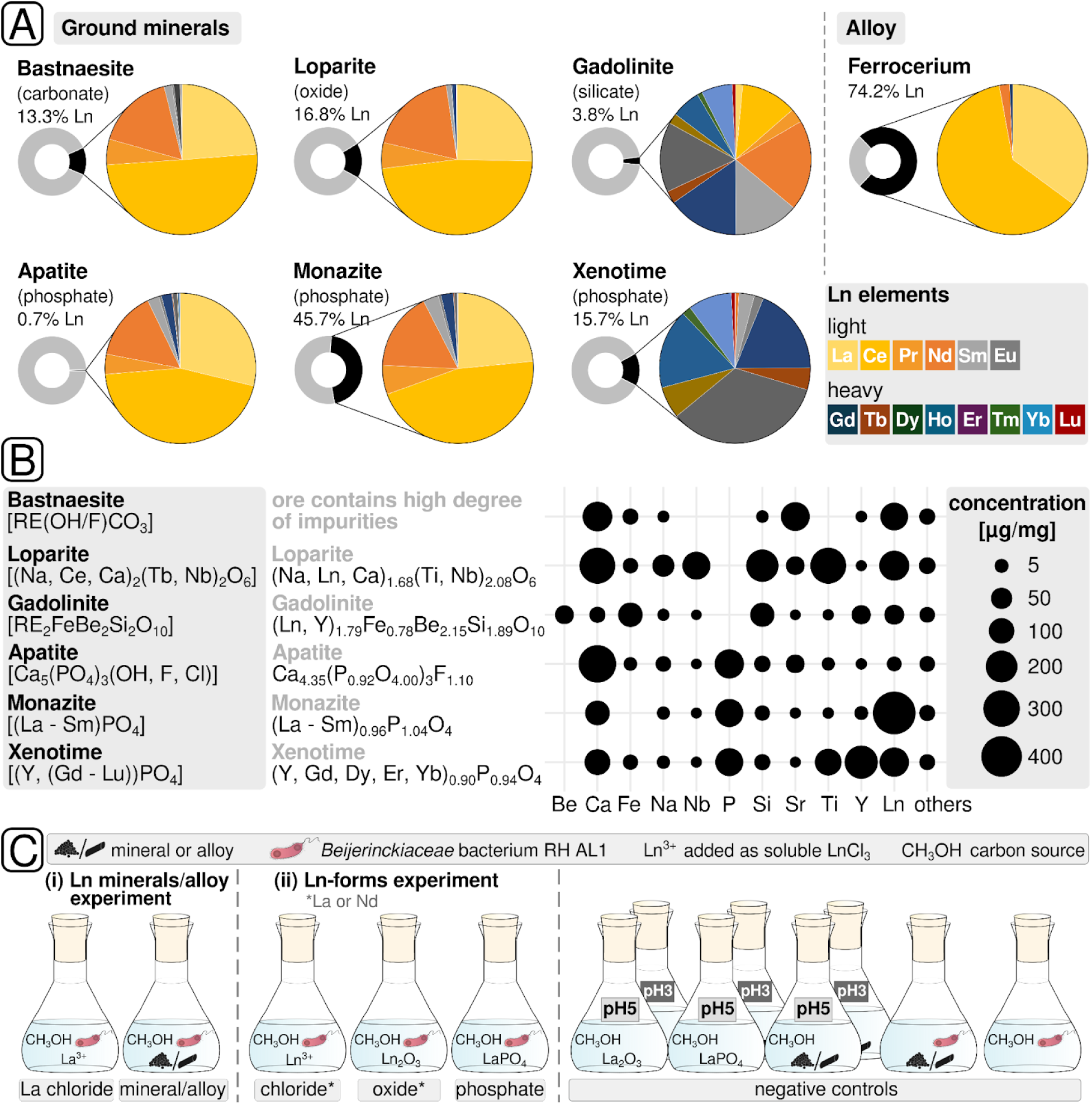
Mineral characteristics and incubation setups. Lns present in the investigated minerals and the alloy ferrocerium were determined via ICP-MS (A), and sum formulas [82–86] and element ratios deduced (B) based on EDX and ICP-MS measurements. Concentrations of detected elements measured via ICP-MS are shown as well. More detailed information about mineral/alloy compositions can be found in the **supplementary material** (**Table S3+4**). In two growth experiments, the effects of (i) Ln-containing minerals or the alloy ferrocerium and (ii) Ln forms (chloride vs. oxide vs. phosphate) were tested (C). Incubations were done in triplicates. Abiotic controls were prepared in two sets with MM2 medium adjusted to pH 5 or pH 3 (reflecting the initial pH used for cultivation and the pH drop observed during cultivation). RE = Rare Earth (**Table S1**).

### Methylotrophic growth in response to different Ln sources

The ability of strain AL1 to utilize different Ln sources for growth on methanol (0.5% [v/v], 123 mM) was tested in two cultivation experiments (Figure 1C, **Table S5**). All tested Ln sources facilitated growth (Figure 2A**+B**). LaCl_3_, Ln_2_O_3_, and loparite showed similar growth patterns, with growth rates (µ) ranging between 0.028 ± 0.004 h^−1^ (LaCl_3_) and 0.035 ± 0.004 h^−1^ (La_2_O_3_). Gadolinite addition led to a slightly lower growth rate (µ = 0.022 ± 0.002 h^−1^) and lower optical densities (OD_600nm_ 0.653 ± 0.019) compared to soluble LaCl_3_ (OD_600nm_ 0.761 ± 0.048). Except for apatite (µ = 0.028 ± 0.005 h^−1^), we observed lower growth rates for cultures supplied with Ln phosphates. Apatite cultures became stationary early at an OD_600nm_ of 0.26 ± 0.02. The highest growth rate (µ = 0.036 ± 0.005 h^−1^) and final optical density (OD_600nm_ 1.324 ± 0.063) were observed with ferrocerium. Abiotic controls supplied with ferrocerium slightly increased in optical density, which was attributed to a reaction between the alloy and the medium.

**Figure 2.**
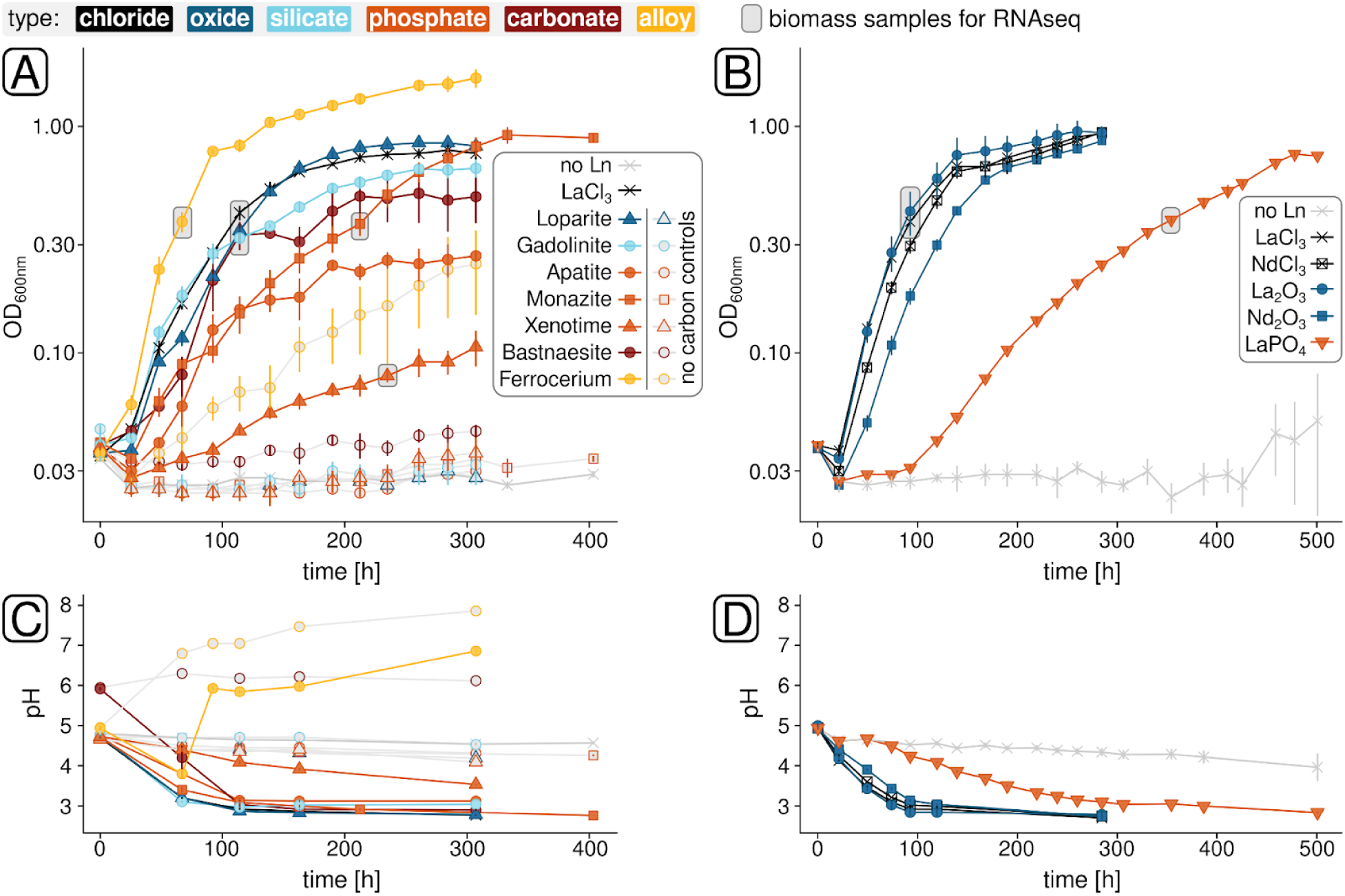
Methylotrophic growth of *Beijerinckiaceae* bacterium RH AL1 with the different tested Ln sources. Growth was either assessed based on incubations with different Ln minerals, and ferrocerium (A), or different forms of La and Nd (B) as Ln source. In addition to optical density measurements, we monitored the pH over the course of the incubations (C + D). Grey boxes highlight the time points at which biomass samples for RNAseq, and supernatant samples for analytics were collected. OD = optical density.

All cultures, except those supplemented with bastnaesite, had an initial pH of 4.7 - 5.0. During exponential growth, the pH decreased to 2.8-3.1 until the cultures entered the stationary phase (Figure 2C**+D**). Incubations with ferrocerium were characterized by a decrease in pH during the first three days of cultivation, followed by an increase accompanied by a color change to orange-brown between 67 and 92 hours of cultivation. Abiotic controls showed a stable pH over time, except for ferrocerium.

### (A)biotic dissolution and intracellular Ln accumulation

Incubations without strain AL1 at pH 5 and pH 3 facilitated the assessment of abiotic Ln dissolution (Figure 3, **Table S7**). We detected less than 1.42 × 10^−2^ µM Lns in any liquid sample at pH 5. Aside from bastnaesite samples, which contained no detectable Ln, the lowest Ln concentrations were detected in ferrocerium (4.74 × 10^−4^ µM Ln) incubations. At pH 3, the minerals were more prone to dissolution. The highest Ln concentrations were detected in samples from gadolinite (3.35 µM Ln) and apatite (1.11 µM Ln) incubations. Detected Ln concentrations were substantially lower for ferrocerium (5.36 × 10^−4^ µM Ln) and the Ln phosphates monazite and xenotime (≤ 1.27 × 10^−1^ µM Ln). Spent media from AL1 cultures tended to show lower Ln concentrations (≤ 2.46 × 10^−2^ µM Ln), except for gadolinite (14.98 µM Ln) and ferrocerium (9.56 µM Ln). Unlike abiotic controls, samples from cultures grown with loparite or phosphate minerals were depleted in light Ln elements.

**Figure 3.**
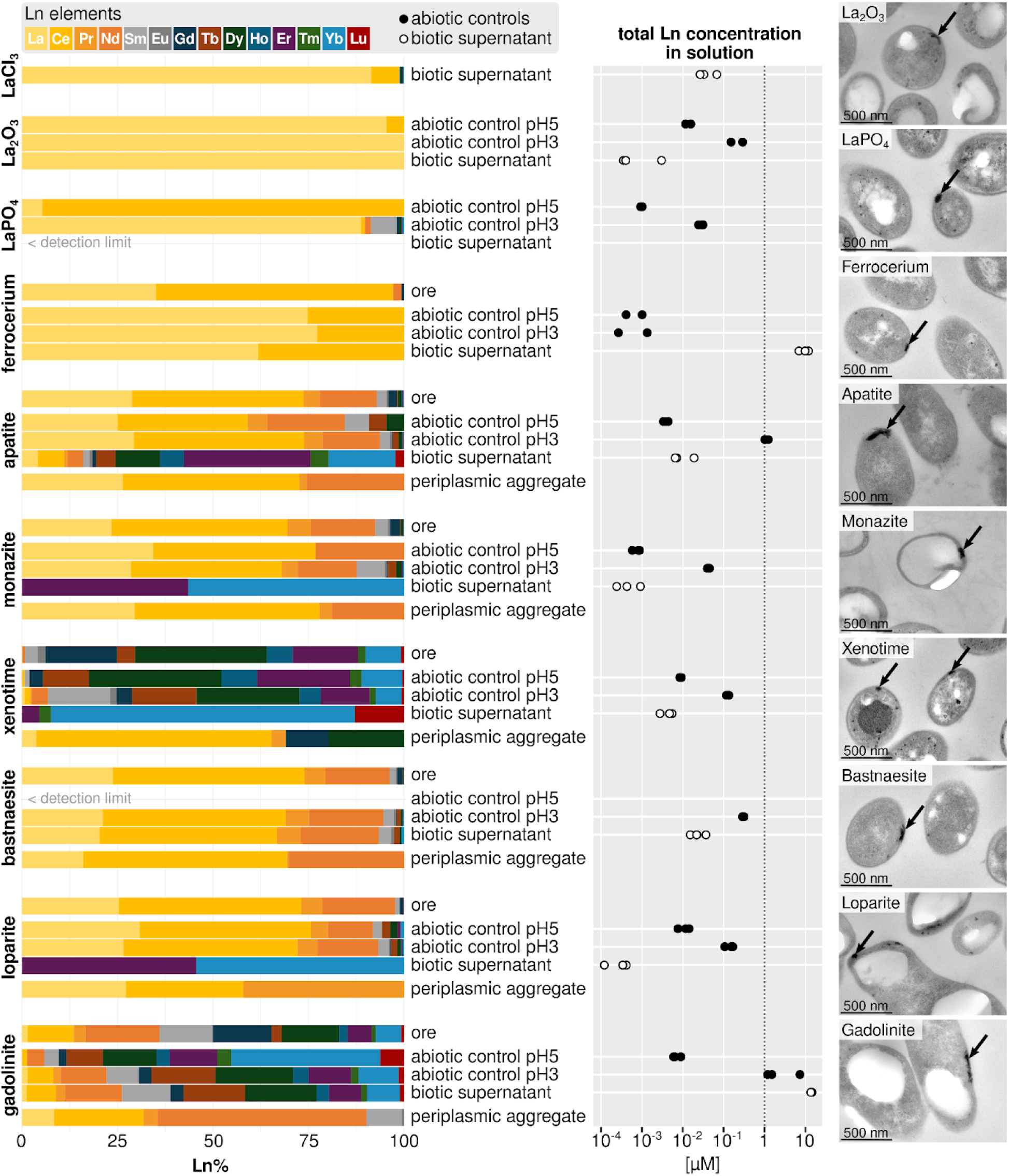
Intracellular Ln accumulation and (a)biotic dissolution. Intracellular Ln accumulation was verified through transmission electron microscopy (TEM). Periplasmic Ln accumulations are highlighted through black arrows. The abundance of Lns in (a)biotic supernatants and periplasmic deposits has been determined by ICP-MS and EDX, respectively. Ln proportions in the Ln sources are the same as in **Figure 1A**. The molarity was deduced from the ICP-MS data and the used cultivation volume. The previously determined optimum concentration for growth with soluble Ln chlorides (1 µM La [25]) is highlighted for orientation. Supernatant analytics was performed in biological and technical replicates. EDX analysis was based on technical triplicates, the TEM micrographs show representative periplasmic deposits.

*Beijerinckiaceae* bacterium RH AL1 has previously been shown to form periplasmic Ln deposits when cultivated with soluble Ln chlorides [5, 27]. Consistent with this, we observed periplasmic deposits in cultivations with the selected Ln sources by TEM (Figure 3). EDX confirmed them to be mineral-like Ln accumulations (**Figure S1+S2, Table S6**). Even cells grown with minerals containing little Lns overall or with a Ln content dominated by heavy Lns showed primarily light Lns in biominerals (Figure 3). Detected element ratios indicated that Lns are accumulated and stored as Ln phosphates.

### Differential gene expression in response to different Ln sources

We used transcriptome-wide gene expression analysis to examine the effects of the Ln sources, focusing on mechanisms for Ln mobilization and differentiation. Differential gene expression analysis was performed based on biological triplicates (**Figures S4 to S6, Table S8+S9**). The ferrocerium incubations yielded only one replicate of sufficient quality for downstream processing. The high iron content likely impaired RNA extraction. Supplementation with different forms of La led to differential gene expression of up to 39% (LaCl_3_ vs. LaO_4_P, 1685 DEGs) and 32% (La_2_O_3_ vs. LaO_4_P, 1392 DEGs) of all encoded genes (Figure 4A**, Table S10+S11**), with substantial overlap of DEGs (n = 1301 genes) between the comparisons. Transcriptome analysis of samples cultivated with different Ln-containing minerals or ferrocerium revealed that between 14% (LaCl_3_ vs. loparite, 599 DEGs) and 53% (monazite vs. ferrocerium, 2298 DEGs) of the encoded genes were differentially expressed. Clustering data sets using Spearman distances, calculated from scaled CPM (counts per million) values, revealed a distinct grouping of Ln phosphates for both carried-out incubation experiments (Figure 4B).

**Figure 4.**
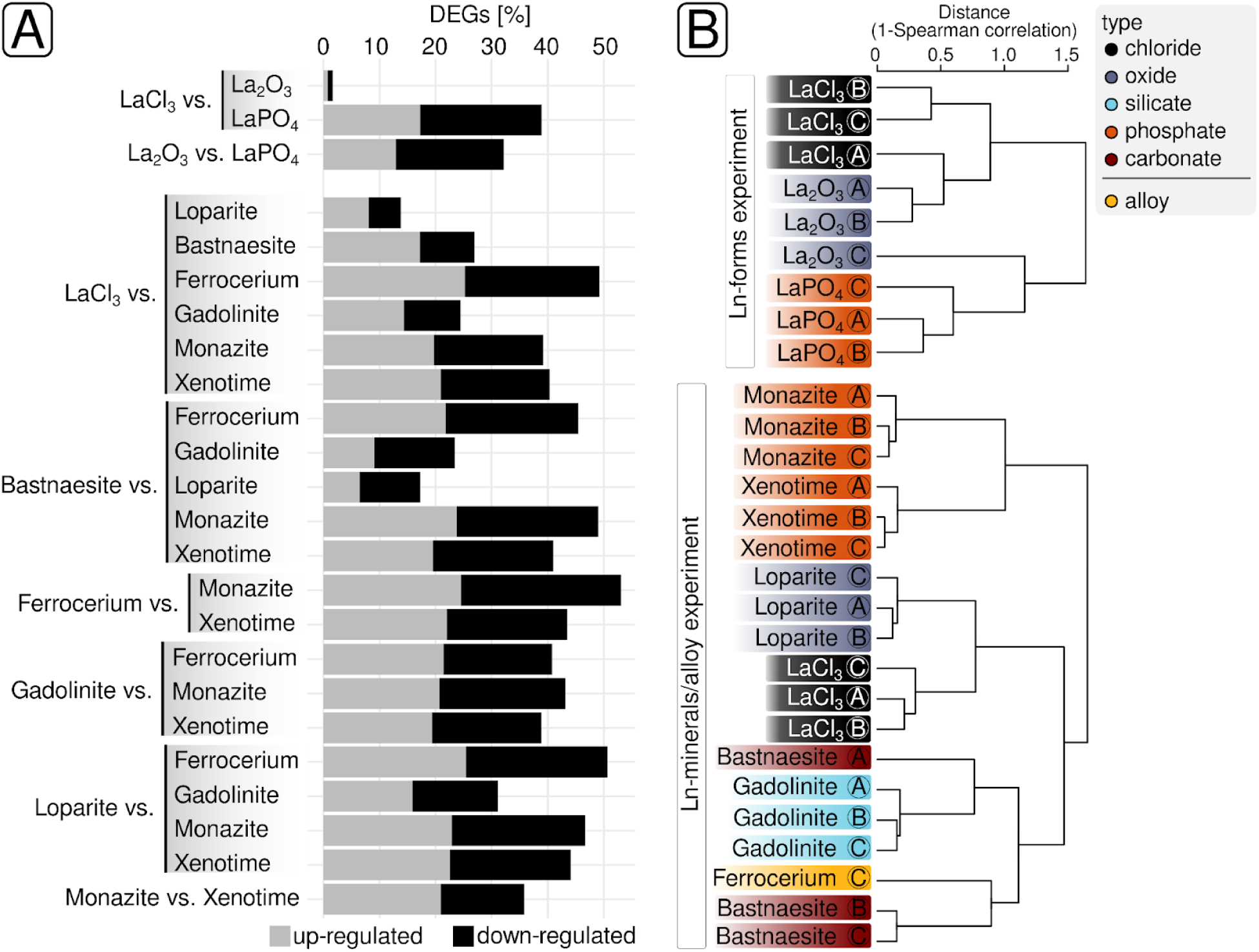
Overview of RNAseq data sets. Statistics regarding up- and downregulated genes from differential gene expression analysis are given (A). Data sets from selected samples were clustered based on Spearman distances (B). Each condition was represented by triplicates, except ferrocerium, due to the high iron load interfering with RNA extraction. Percentages are calculated based on the total number of genes in the genome of the *Beijerinckiaceae* bacterium RH AL1. DEGs = differentially expressed genes.

### Gene expression dynamics of the lanthanome

A closer look at lanthanome genes (Figure 5, **Figure S7**, **Table S12+S13**) revealed that especially Ln phosphates (LaPO_4_, monazite, xenotime) caused notable differences in gene expression (Figure 5). These were particularly evident when looking at *lanM* (RHAL1_01396, encoding the periplasmic Ln-binding protein lanmodulin), *lanD* (RHAL1_03683, coding for landiscernin, a protein supposed to interact with LanM), *lutH* (RHAL1_00153), and *xoxJ* (RHAL1_02996). Compared to LaCl_3_, La_2_O_3_, and non-phosphate Ln minerals, *lanM* and *lanD* were downregulated (log_2_FC *lanM* between −2.05 and −3.75, log_2_FC *lanD* between −1.49 and −2.83). The *lutH* gene (coding for a TonB-dependent receptor involved in periplasmic Ln uptake) was upregulated, reaching log_2_CPM values between 10.32 and 11.19 in samples originating from incubations with Ln phosphates (Figure 5, **Figure S7**). The *xoxJ* gene (RHAL1_02996, XoxJ is assumed to be involved in XoxF activation) showed broad expression ranging from 4.41 to 8.22 log_2_CPM (**Table S12+S13**) and was strongly upregulated when comparing phosphates to LaCl_3_ (log_2_FC 2.91 - 3.58) (**Figure S7**).

**Figure 5.**
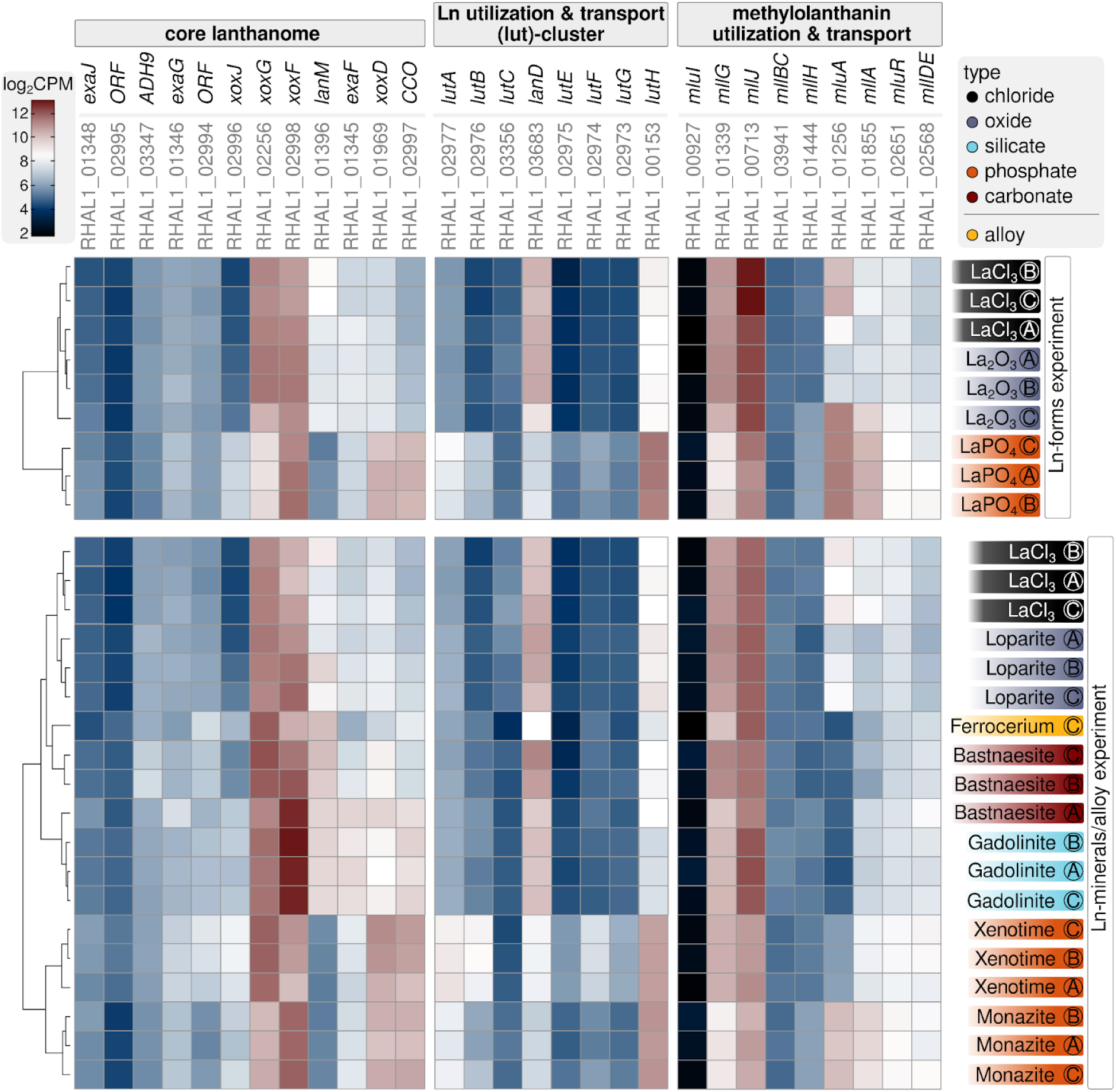
Lanthanome-related gene expression dynamics. The focus was on core lanthanome genes, genes directly linked to Ln-driven methylotrophy/alcohol oxidation, as well as gene homologs of the *lut*( lanthanide <u>u</u>tilization and transport)- and *mll/mlu* (<u>m</u>ethylolanthanin/<u>m</u>ethylolanthanin <u>u</u>ptake)-clusters. Gene expression is given in log_2_CPM (log_2_ <u>c</u>ounts <u>p</u>er <u>m</u>illion). ORF = open reading frame, CCO = cytochrome C oxidase, ADH9 = gene encoding an Ln-dependent alcohol dehydrogenase that belongs to subclade 9 within the PQQ ADH family.

The expression of *xoxF* and *xoxG*, coding for the Ln-dependent methanol dehydrogenase and its physiological electron acceptor, a c_L_-type cytochrome, central to Ln utilization in AL1, was overall high (*xoxF* log_2_CPM 9.72 - 13.03, *xoxG* 9.11 - 11.88). This was independent of the supplied Ln source. The gene expression ratio between these two genes was near one and only shifted slightly depending on the Ln source (**Figure S8**). When Ln phosphates were omitted, multiple genes associated with the *lut*-cluster were found to be nonresponsive to differences in lanthanide supplementation. This also included *lutAEF*, which encode the periplasmic binding protein (*lutA*), ATP-binding cassette (*lutE*), and transmembrane component (*lutF*) of an ABC transporter involved in Ln import from the peri- into the cytoplasm. Gene homologues of the *mll-*/*mlu-cluster* partially responded to different Ln sources. Compared to LaCl_3_, *mluA* (RHAL1_01256, encoding a TonB-dependent receptor) was downregulated when AL1 was grown with ferrocerium, bastnaesite, and gadolinite (log_2_FC between −3.66 and −4.01). We also noted differences in gene expression responses, dependent on the different Ln phosphates.

### Identification of clusters of correlated genes

Network analysis based on weighted gene co-expression identified 9 modules comprising between 60 and 1015 genes. Gene expression data were summarized module-wise based on eigengenes (ME-1 through ME-9) (Figure 6A**+B, Figures S9 to S11**, **Table S14**). The response to different Ln phosphates varied, particularly among genes assigned to modules 1-3 and 8. Correlation analysis showed a positive link between individual eigengenes (ME-1 to ME-3) and different Ln phosphates (Figure 6C). We also observed a positive correlation between ME-7 and the silicate gadolinite. Selected modules were examined, focusing on genes with high module membership (MM, the correlation between individual module genes and the respective eigengene) and strong correlations with Ln sources (**Figure S12+S13**). We were especially interested in genes putatively associated with Ln mobilization, uptake, storage and discrimination.

**Figure 6.**
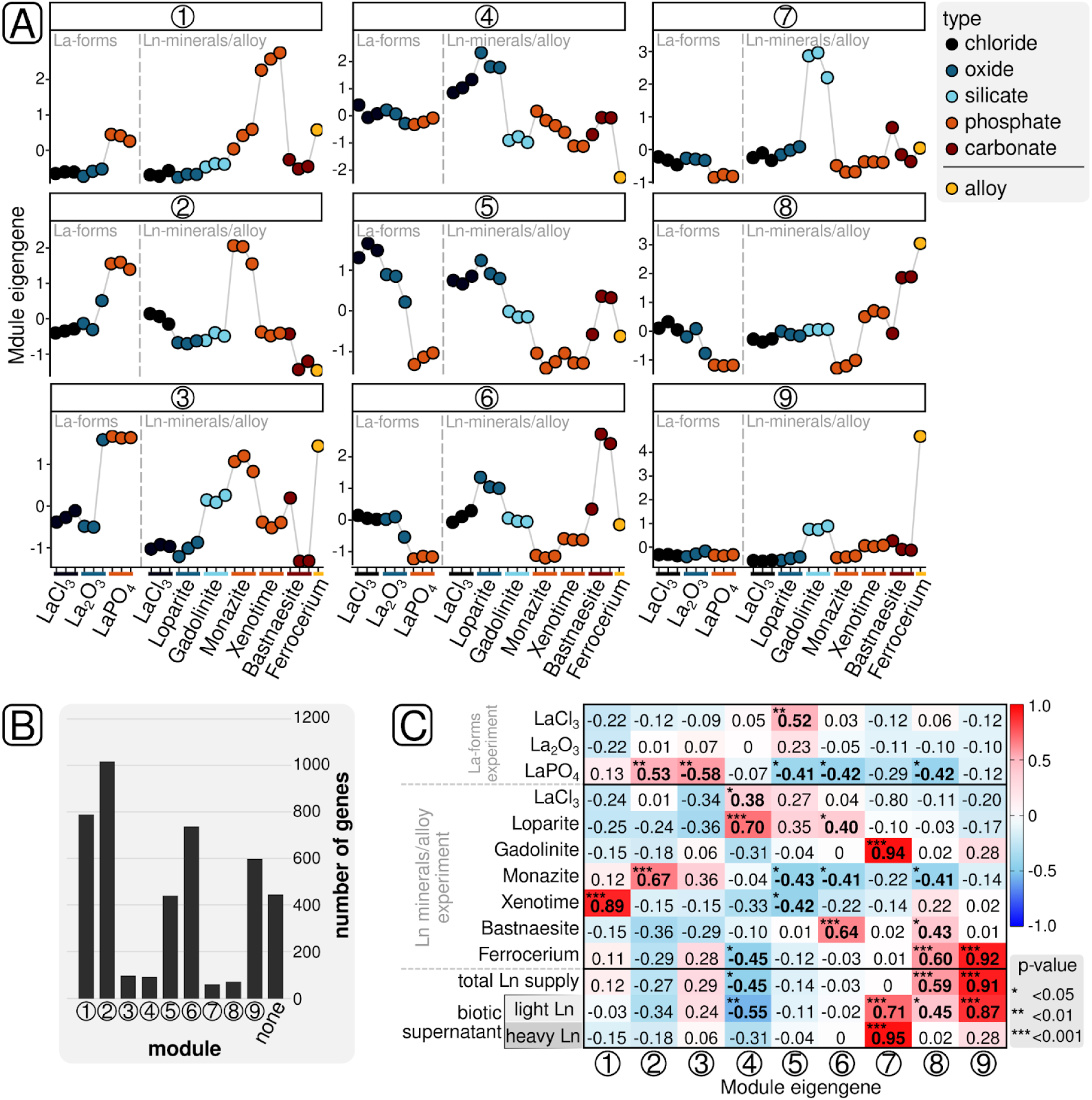
Gene correlation network analysis. Eigengene expression profiles have been deduced from modules obtained via weighted gene co-expression network analysis (WGCNA). WGCNA led to the identification of nine modules, and the expression profiles of respective module eigengenes are shown (A). Sample colors reflect the respective type of Ln source, which was supplied for cultivation. Modules comprised between 60 and 1015 genes (B). Pearson correlations between module eigengenes and selected traits (Ln sources, total Ln supply, and Ln content in biotic supernatant) have been calculated (C). Different significance levels are highlighted.

Module 1 contained several genes associated with PQQ biosynthesis and the lanthanome (Figure 7). We observed particularly high MM for three *pqqA* copies (RHAL1_01618, RHAL1_02672, RHAL1_02989, coding for PqqA, the precursor peptide for PQQ synthesis) and *lut*-cluster genes *lutABEFG* (RHAL1_02977 - RHAL1_02973) (MM between 0.857 and 0.972, p < 0.001). The genes *lutH* and *mluA* (RHAL1_01256), encoding TonB-dependent receptors, were grouped into module 2 (MM 0.765 and 0.719, p < 0.001). A closer look at this module revealed various genes associated with iron homeostasis, including uptake (RHAL1_p00067, high-affinity iron transporter; RHAL1_03444-RHAL1_03446, siderophore-independent Fe uptake system EfeUOB) and storage (RHAL1_00987, bacterioferritin) (MM between 0.741 and 0.901, p < 0.001). Module 2 also contained genes *pstSABC* and *phoUB* (RHAL1_00909 - RHAL1_00914) associated with phosphate transport (MM between 0.708 and 0.878, p < 0.001). Genes *xoxF* (RHAL1_02998) and *exaF* (RHAL1_01345) displayed distinct differences in expression profiles compared to other lanthanome-associated genes and were grouped in module 7 (MM 0.861 and 0.811, p < 0.001). Module 6 comprised multiple genes associated with redox stress and balance, including genes *ssuABCD* (RHAL1_02408 - RHAL1_02411, MM between 0.826 and 0.917, p < 0.001) linked to alkanesulfonate uptake. Several chaperone encoding genes, including *groEL* and *groES,* showed high memberships for module 2, as well as genes encoding glutaredoxin and thioredoxin. Both *lanM* (RHAL1_01396, MM = 0.696 with p < 0.001) and *lanD* (RHAL1_03683, MM = 0.871 with p < 0.001) were part of module 6, but MM was lower for *lanM*.

**Figure 7.**
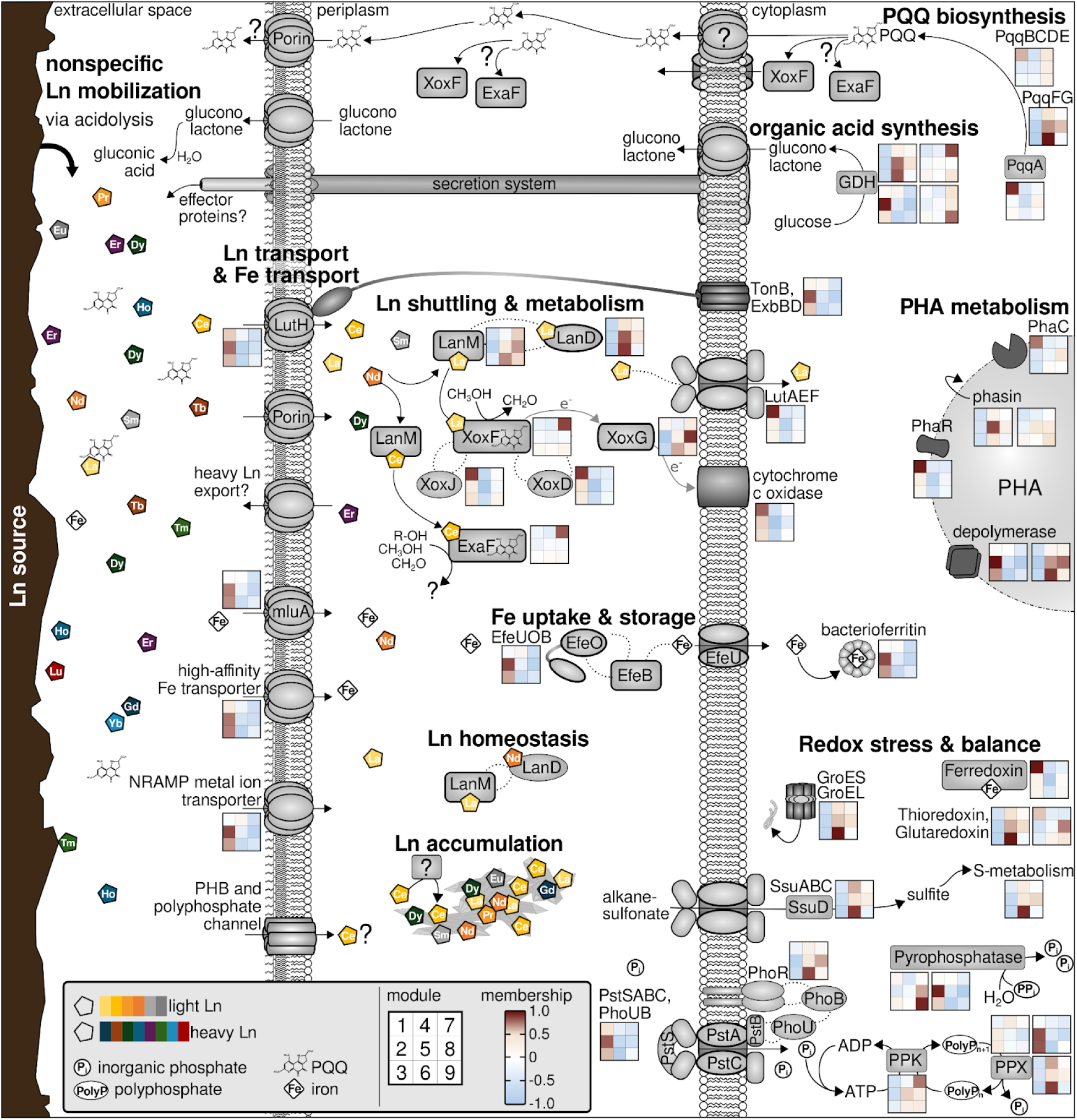
Physiological response to changes in Ln supplementation based on module membership. The individual heatmaps indicate the module membership of the genes encoding the shown proteins and groups of proteins, which were selected based on module memberships, and correlations with Ln sources. The focus was on proteins assumed to be involved in Ln mobilization, uptake, shuttling, and discrimination. Parts of the lanthanome are visualized, including Ln-dependent ADHs as key enzymes for methanol oxidation and Ln-binding proteins involved in Ln shuttling and potentially homeostasis. In addition, module memberships of genes encoding proteins involved in Polyhydroxyalkanoate (PHA) biosynthesis and degradation are shown, as well as cellular mechanisms involved in redox stress and balance. GDH = glucose dehydrogenase, PPK = polyphosphate kinase, PPX = exopolyphosphatase.

The genome of AL1 contains four glucose dehydrogenase genes (RHAL1_00125, RHAL1_01212, RHAL1_02698, and RHAL1_3913), which are linked to gluconolactone (a potential metal chelator) formation. All four of these genes clustered into different modules (modules 6 through 9, MM between 0.765 and 0.947, p < 0.001) (**Table S14**).

## DISCUSSION

We could show that its metabolic flexibility equips *Beijerinckiaceae* bacterium RH AL1 to mobilize, selectively take up, and store utilizable Lns from diverse sources that differ in type and Ln content. The recalcitrance and the availability of light, utilizable Lns, but not the overall Ln content, controlled Ln-dependent, methylotrophic growth. Ln-utilization is centered around Ln-dependent PQQ ADHs, which are, despite intense research over the last years, still the only known class of Ln-dependent enzymes [46, 47]. The incorporation of heavy Lns as cofactors in Ln-dependent PQQ ADH must be avoided by a discrimination mechanism during Ln handling. Ln-utilizing microbes have a preference for light Lns [18], which is rooted in PQQ ADHs being paired with c_L_-type cytochromes as physiological electron acceptors. These are tuned in terms of redox potential to PQQ ADH incorporating light Ln [48]. Heavy Lns can impair PQQ ADH activity through differences in Lewis acidity and size, the latter potentially perturbing the active site [49, 50].

Acidolysis, the dissolution of solids by decreasing the pH, forms the backbone of Ln mobilization in AL1. Based on our gene expression data, it is driven by the release of organic acids such as gluconolactone (GL), which is well known for phosphate-solubilizing bacteria that are also studied for metal- and Ln-leaching [51–53]. The AL1 genome encodes four glucose dehydrogenases, potentially involved in GL formation. Gene correlation network analysis suggested fine-tuned expression depending on the supplemented Ln source.

Acidolysis alone confers no selectivity. The analysis of (a)biotic supernatants revealed the preferential depletion of light Ln, indicating Ln discrimination during uptake. Organic acids such as citric acid, oxalic acid, and GL are potent metal chelators with Ln-binding preferences based on differences in Lewis acidity, ionic radii, and coordination geometry [54–57]. Besides its role as a cofactor in PQQ ADHs, PQQ was shown, outside of a protein context, to form complexes with preferentially light Lns [58, 59]. Differential expression of AL1’s *pqqA* genes, encoding the PQQ precursor peptide, suggests a role for PQQ in Ln mobilization and uptake, especially with respect to Ln phosphates. The availability and interplay between different chelators can aid AL1 discriminating Lns during mobilization and uptake.

The identification of the *lut-* and *mll*-/*mlu*-clusters [11, 13] provided evidence that Ln uptake by mesophilic bacteria is metallophore-based, similar to iron and copper uptake, and centered around TonB-dependent receptors and ABC transporters. Methylolanthanin is the first characterized Ln-binding metallophore [13, 60], and related metallophores have been proposed for *M. aquaticum* 22A [61]. Siderophores and chalkophores have distinct metal-binding preferences [62], conveying selectivity at the stage of mobilization. The binding preferences of methylolanthanin regarding Lns are inconclusive [60]. The need for metal-binding metallophores depends on bioavailability. Organisms such as *Methylacidiphilum fumariolicum* SolV, thriving in acidic systems with high metal loads, have no need for metallophore-based uptake [63]. Strain AL1 is likewise less reliant on metal chelators for accessing Lns, as it is a mild acidophile [5, 25].

Gene homologs of the *lut*- and *mll*-/*mlu*-clusters have been identified in AL1, but the observed amino acid sequence identities are partially rather low [5, 25]. This dissimilarity was reflected in the nonresponsiveness of some of these genes to changes in Ln supplementation. Notable exceptions were *lutH* and *mluA*, which encode TonB-dependent receptors within the *lut*- and *mll*-/*mlu*-clusters. The substrates for periplasmic uptake by these receptors are unknown in any organism. Gene expression dynamics and network analysis indicated that LutH may be linked to Ln sensing, while MluA appears to bridge Ln and iron metabolism via EfeUOB and bacterioferritin. EfeUOB is a siderophore-independent iron uptake system [64, 65]. EfeO was found to bind Sm^3+^ in a recent study [66], and *efeUOB* were differentially expressed in strain AL1 when swapping LaCl_3_ for an equimolar Ln cocktail [5]. Crosstalk between Ln and iron metabolism was suggested for *Pseudomonas alloputida* KT2440 based on overall metal availability [67] and supported by observed effects of iron levels on the concentrations of methylolanthanin and cell-associated Nd in *M. extorquens* AM1 cultivations [60]. From studies of iron homeostasis, it is known that TonB-ABC transport systems are downregulated in response to iron availability [68, 69].

We previously proposed that periplasmic Ln deposition is key to Ln homeostasis [5, 27]. AL1 enriched light, utilizable Ln as mineral-like periplasmic deposits from all tested sources, despite in part poor solubilities, an overall low Ln content, and low proportions of utilizable Lns. The underrepresentation of heavy Lns in the mineral-like deposits, together with their overrepresentation in spent medium, supports that Ln discrimination occurs during uptake. The low proportions of heavy Lns taken up are diverted into periplasmic deposits, preventing downstream incorporation during protein maturation.

As shown before [5, 27], periplasmic accumulation occurred at the cell poles, in close proximity to PHB granules. Channels composed of complexed short-chain PHB (polyhydroxybutyrate, a common form of PHA for carbon storage) and polyphosphate represent one important uptake route for calcium [70–73]. Correlating gene expression patterns of selected lanthanome- and PHA-related genes supports a link between PHA and Ln metabolism, but this warrants further investigation. We previously hypothesized [5] that the involvement of such channels in Ln uptake can account for the localisation of the periplasmic deposits and contribute to selective Ln uptake.

Ln discrimination during Ln uptake is further supported by examining the core components of the lanthanome. XoxF tends not to incorporate heavy Lns *in vivo [74]*, and if they are bound, activity is strongly reduced [50]. Heavy Lns impair the redox cycling of reduced PQQ [75]. Besides, XoxF mismetallation with heavy Lns interferes with methanol oxidation by distorting the electron transfer from XoxF to its physiological electron acceptor, XoxG. The expression of *xoxF* in AL1 differed between the Ln sources, likely reflecting adaptation and the need for flexibility to changing Ln availability. However, the expression ratio between *xoxF* and *xoxG*, which we considered as a proxy for the misincorporation of heavy Lns into XoxF, was stable, reflecting that misincorporation, if happening at all, was negligible. Ln discrimination in AL1 occurs upstream of PQQ ADH maturation.

Recent work described an intriguing, siphon-like mechanism for sorting Lns, built around lanmodulin (LanM) and landiscerin (LanD) in *M. extorquens* AM1 [10]. Heavy Lns are prevented from cytoplasmic uptake and downstream incorporation into PQQ ADHs by transferring them to LanM through LanD. The Ln binding preferences of LanM and LanD match, and LanD is equipped with a novel Ln binding motif, distinct from the heavily studied EF-hand motif of LanM. We previously showed that *lanM* and *lanD* homologs in AL1 are upregulated when the strain is grown with a mixture of Lns rather than La, supplied as soluble chlorides, supporting their involvement in periplasmic Ln sorting [5]. Here, we observed downregulation of *lanM* and *lanD* when AL1 was grown with chemically stable Ln phosphates, including xenotime, which has an elevated load of heavy Lns, likely limiting overall Ln influx. LanM and LanD arguably contribute to the periplasmic Ln balance in AL1, but their roles in Ln discrimination remain open questions. The encoded LanD homolog in AL1 differs at the amino acid level with respect to the predicted Ln-binding site.

The assembled gene expression data provide further evidence that Lns reach into many aspects of cellular metabolism. In organisms featuring the Ln switch, gene expression dynamics tend to be limited to the lanthanome and methylotrophy-related genes [6, 12, 76–78]. The Ln switch is rather the exception, and organisms without it are more common in the environment [46, 79–81]. The lack of gene expression data from bacteria lacking the Ln switch is a blind spot that needs to be addressed in the future. We showed that AL1 gene expression was primarily affected by the type of Ln source. A high number of differentially expressed genes indicated a complex metabolic response to the various Ln sources, underscoring the need to use more natural Ln sources when studying microbial Ln utilization. The mineral type represented the predominant factor driving gene expression changes. Many of the genes we found to be differentially expressed were previously identified by us in our work with Ln chloride salts [5]. These included, among others, genes associated with chemotaxis and motility, polyhydroxyalkanoates, and sulfur metabolism. In addition to Ln element concentrations and proportions, natural Ln sources also differed in their concentrations of trace elements such as iron, which are essential for cellular metabolism. The effects we have only been able to classify so far by examining different La forms and based on previous studies with different Ln elements and La concentrations [5] could be observed more precisely in the future through the use of synthetic minerals.

Taken together, our data suggest that Ln discrimination in *Beijerinckiaceae* bacterium RH AL1 occurs through a multilayered process. Acidolysis provides access to mineral-bound lanthanides, but is intrinsically non-selective. Selectivity only arises through the interplay of chelation, different uptake mechanisms, and periplasmic storage, which facilitate the preferential enrichment of utilizable light Lns while heavy Lns are diverted. Ln discrimination in AL1 is sequential with potential implications for bioinspired Ln recovery strategies.

## ACKNOWLEDGEMENTS

The authors are grateful to the late Herfried Richter for samples of bastnaesite, loparite, gadolinite, and apatite, and to the Treibacher Industrie AG (Althofen, Austria) for providing the used ferrocerium. The authors are grateful to Dirk Merten for organizing the ICP-MS measurements of (a)biotic supernatant samples. LG acknowledges Melissa Tuere’s help with cultivation. The authors thank the employees of the central analytic facilities at Ludwig Maximilians University Munich (Munich, Germany). Sequencing was carried out by the sequencing core facility of the Leibniz Institute on Aging-Fritz Lipmann Institute (Jena, Germany). This research was supported by the Deutsche Forschungsgemeinschaft (DFG) (project number: 467713456, grant: WE6579/4-1; granted to CEW), through the SFB 1127 ChemBioSys (project number 239748522), and by the de.NBI Cloud within the German Network for Bioinformatics Infrastructure (de.NBI) and ELIXIR-DE (Forschungszentrum Jülich and W-de.NBI-001, W-de.NBI-004, W-de.NBI-008, W-de.NBI-010, W-de.NBI-013, W-de.NBI-014, W-de.NBI-016, W-de.NBI-022) as well as the ERC Starting Grant Lanthanophore. LG thanks the Jena School for Microbial Communication (JSMC) for its financial support.

## AUTHOR CONTRIBUTIONS

Linda Gorniak and Carl-Eric Wegner conceived this study. Linda Gorniak was responsible for cultivation and molecular work. Linda Gorniak and Carl-Eric Wegner carried out RNAseq data analysis. Sophie M. Gutenthaler-Tietze, Alina Lobe, Robin Steudtner, and Lena J. Daumann characterized the Ln sources used. Supernatant analytics was provided by Thorsten Schäfer. Martin Westermann and Frank Steiniger conducted advanced electron microscopy of periplasmic Ln accumulations. Lena J. Daumann, Kirsten Küsel, and Thorsten Schäfer provided resources. Carl-Eric Wegner secured funding and was responsible for project administration and main supervision. Linda Gorniak was responsible for data visualization and wrote the first draft with edits from Carl-Eric Wegner. All authors read and contributed to the final version of the manuscript

## REFERENCES

1. Rare earths in the energy transition: what threats are there for the ‘vitamins of modern society’? IFPEN. https://www.ifpenergiesnouvelles.com/article/les-terres-rares-transition-energetique-quelles-menaces-les-vitamines-lere-moderne. Accessed 1 Apr 2022.

2. European Commission: Directorate-General for Internal Market, Industry, Entrepreneurship and SMEs, Study on the critical raw materials for the EU 2023 – Final report, Publications Office of the European Union, 2023, https://data.europa.eu/doi/10.2873/725585

3. Wehrmann M, Billard P, Martin-Meriadec A, Zegeye A, Klebensberger J. Functional role of lanthanides in enzymatic activity and transcriptional regulation of pyrroloquinoline quinone-dependent alcohol dehydrogenases in *Pseudomonas putida* KT2440. MBio 2017; 8: mbio.00570-17.

4. Wehrmann M, Toussaint M, Pfannstiel J, Billard P, Klebensberger J. The cellular response to lanthanum is substrate specific and reveals a novel route for glycerol metabolism in *Pseudomonas putida* KT2440. MBio 2020; 11: mbio.00516-20.

5. Gorniak L, Bechwar J, Westermann M, Steiniger F, Wegner C-E. Different lanthanide elements induce strong gene expression changes in a lanthanide-accumulating methylotroph. Microbiol Spectr 2023; 11: e0086723.

6. Gorniak L, Bucka SL, Nasr B, Cao J, Hellmann S, Schäfer T, et al. Changes in growth, lanthanide binding, and gene expression in *Pseudomonas alloputida* KT2440 in response to light and heavy lanthanides. mSphere 2024; 9: e0068524.

7. Mattocks JA, Ho JV, Cotruvo JA Jr. A Selective, Protein-Based Fluorescent Sensor with Picomolar Affinity for Rare Earth Elements. J Am Chem Soc 2019; 141: 2857–2861.

8. Cotruvo JA Jr, Featherston ER, Mattocks JA, Ho JV, Laremore TN. Lanmodulin: A Highly Selective Lanthanide-Binding Protein from a Lanthanide-Utilizing Bacterium. J Am Chem Soc 2018; 140: 15056–15061.

9. Hemmann JL, Keller P, Hemmerle L, Vonderach T, Ochsner AM, Bortfeld-Miller M, et al. Lanpepsy is a novel lanthanide-binding protein involved in the lanthanide response of the obligate methylotroph *Methylobacillus flagellatus*. J Biol Chem 2023; 299: 102940.

10. Larrinaga WB, Jung JJ, Lin C-Y, Boal AK, Cotruvo JA Jr. Modulating metal-centered dimerization of a lanthanide chaperone protein for separation of light lanthanides. Proc Natl Acad Sci U S A 2024; 121: e2410926121.

11. Roszczenko-Jasińska P, Vu HN, Subuyuj GA, Crisostomo RV, Cai J, Lien NF, et al. Gene products and processes contributing to lanthanide homeostasis and methanol metabolism in *Methylorubrum extorquens* AM1. Sci Rep 2020; 10: 12663.

12. Ochsner AM, Hemmerle L, Vonderach T, Nüssli R, Bortfeld-Miller M, Hattendorf B, et al. Use of rare-earth elements in the phyllosphere colonizer *Methylobacterium extorquens* PA1. Mol Microbiol 2019; 111: 1152–1166.

13. Zytnick AM, Gutenthaler-Tietze SM, Aron AT, Reitz ZL, Phi MT, Good NM, et al. Identification and characterization of a small-molecule metallophore involved in lanthanide metabolism. Proc Natl Acad Sci U S A 2024; 121: e2322096121.

14. Dong Z, Mattocks JA, Deblonde GJ-P, Hu D, Jiao Y, Cotruvo JA Jr, et al. Bridging Hydrometallurgy and Biochemistry: A Protein-Based Process for Recovery and Separation of Rare Earth Elements. ACS Cent Sci 2021; 7: 1798–1808.

15. Voutsinos MY, Banfield JF, McClelland H-LO. Extensive and diverse lanthanide-dependent metabolism in the ocean. ISME J 2025; 19: wraf057.

16. Gonzalez V, Vignati DAL, Leyval C, Giamberini L. Environmental fate and ecotoxicity of lanthanides: are they a uniform group beyond chemistry? Environ Int 2014; 71: 148–157.

17. Kotelnikova AD, Rogova OB, Stolbova VV. Lanthanides in the Soil: Routes of Entry, Content, Effect on Plants, and Genotoxicity (a Review). Eurasian Soil Sci 2021; 54: 117–134.

18. Daumann LJ. Essential and ubiquitous: The emergence of lanthanide metallobiochemistry. Angew Chem Int Ed Engl 2019; 58: 12795–12802.

19. Zaharescu DG, Burghelea CI, Dontsova K, Presler JK, Maier RM, Huxman T, et al. Ecosystem Composition Controls the Fate of Rare Earth Elements during Incipient Soil Genesis. Sci Rep 2017; 7: 43208.

20. Banfield JF, Eggleton RA. Apatite Replacement and Rare Earth Mobilization, Fractionation, and Fixation During Weathering. Clays Clay Miner 1989; 37: 113–127.

21. Firsching FH, Brune SN. Solubility products of the trivalent rare-earth phosphates. J Chem Eng Data 1991; 36: 93–95.

22. Taunton AE, Welch SA, Banfield JF. Microbial controls on phosphate and lanthanide distributions during granite weathering and soil formation. Chem Geol 2000; 169: 371–382.

23. U.S. Geological Survey. Mineral commodity summaries 2025. 2025. U.S. Geological Survey.

24. Shi S, Pan J, Dong B, Zhou W, Zhou C. Bioleaching of rare earth elements: Perspectives from mineral characteristics and microbial species. Minerals (Basel) 2023; 13: 1186.

25. Wegner C-E, Gorniak L, Riedel S, Westermann M, Küsel K. Lanthanide-Dependent Methylotrophs of the Family Beijerinckiaceae: Physiological and Genomic Insights. Appl Environ Microbiol 2020; 86: e01830–19.

26. Wegner C-E, Liesack W. Unexpected Dominance of Elusive Acidobacteria in Early Industrial Soft Coal Slags. Front Microbiol 2017; 8: 1–13.

27. Wegner C-E, Westermann M, Steiniger F, Gorniak L, Budhraja R, Adrian L, et al. Extracellular and Intracellular Lanthanide Accumulation in the Methylotrophic *Beijerinckiaceae* Bacterium RH AL1. Appl Environ Microbiol 2021; 87: e0314420.

28. Dedysh SN, Kulichevskaya IS, Serkebaeva YM, Mityaeva MA, Sorokin VV, Suzina NE, et al. *Bryocella elongata* gen. nov., sp. nov., a member of subdivision 1 of the Acidobacteria isolated from a methanotrophic enrichment culture, and emended description of *Edaphobacter aggregans* Koch et al. 2008. Int J Syst Evol Microbiol 2012; 62: 654–664.

29. Dedysh SN, Dunfield PF. 2014. Cultivation of methanotrophs, p 231–247. In McGenity T, Timmis K, Nogales B (ed), Hydrocarbon and lipid microbiology protocols. Springer protocols handbooks. Springer, Berlin, Germany

30. Andrews S. FastQC: a quality control tool for high throughput sequence data. http://www.bioinformatics.babraham.ac.uk/projects/fastqc..

31. R Core Team. R: A Language and Environment for Statistical Computing. 2022. R Foundation for Statistical Computing, Vienna, Austria.

32. Robinson MD, McCarthy DJ, Smyth GK. edgeR: a Bioconductor package for differential expression analysis of digital gene expression data. Bioinformatics 2010; 26: 139–140.

33. Ritchie ME, Phipson B, Wu D, Hu Y, Law CW, Shi W, et al. limma powers differential expression analyses for RNA-sequencing and microarray studies. Nucleic Acids Res 2015; 43: e47.

34. Rohart F, Gautier B, Singh A, Lê Cao K-A. mixOmics: An R package for ‘omics feature selection and multiple data integration. PLoS Comput Biol 2017; 13: e1005752.

35. Rau A, Gallopin M, Celeux G, Jaffrézic F. Data-based filtering for replicated high-throughput transcriptome sequencing experiments. Bioinformatics 2013; 29: 2146–2152.

36. Rutter L, Cook D. bigPint: A Bioconductor visualization package that makes big data pint-sized. PLoS Comput Biol 2020; 16: e1007912.

37. Langfelder P, Horvath S. WGCNA: an R package for weighted correlation network analysis. BMC Bioinformatics 2008; 9: 559.

38. Langfelder P, Horvath S. Fast R functions for robust correlations and hierarchical clustering. J Stat Softw 2012; 46: i11.

39. Wickham H. ggplot2. WIREs Comp Stat 2011; 3: 180–185.

40. Warnes, Bolker, Bonebakker, Gentleman. gplots: various R programming tools for plotting data, version 3.0. 1. Search in.

41. Kassambara A. ggpubr:‘ggplot2’ based publication ready plots (Version 0.1. 7). Obtido desde https://CRANR-project org/package= ggpubr 2018.

42. Wilke CO. cowplot: streamlined plot theme and plot annotations for ‘ggplot2’. R package version 1.0. 0. 2019. See https://CRAN.R-project.org/package=cowplot.

43. Conway JR, Lex A, Gehlenborg N. UpSetR: an R package for the visualization of intersecting sets and their properties. Bioinformatics 2017; 33: 2938–2940.

44. Köster J, Rahmann S. Snakemake—a scalable bioinformatics workflow engine. Bioinformatics 2012; 28: 2520–2522.

45. Mölder F, Jablonski KP, Letcher B, Hall MB, Tomkins-Tinch CH, Sochat V, et al. Sustainable data analysis with Snakemake. F1000Res 2021; 10: 33.

46. Keltjens JT, Pol A, Reimann J, Op Den Camp HJM. PQQ-dependent methanol dehydrogenases: Rare-earth elements make a difference. Appl Microbiol Biotechnol 2014; 98: 6163–6183.

47. Daumann LJ, Pol A, Op den Camp HJM, Martinez-Gomez NC. A perspective on the role of lanthanides in biology: Discovery, open questions and possible applications. Adv Microb Physiol 2022; 81: 1–24.

48. Featherston ER, Rose HR, McBride MJ, Taylor EM, Boal AK, Cotruvo JA Jr. Biochemical and structural characterization of XoxG and XoxJ and their roles in lanthanide-dependent methanol dehydrogenase activity. Chembiochem 2019; 20: 2360–2372.

49. Lumpe H, Pol A, Op den Camp HJM, Daumann LJ. Impact of the lanthanide contraction on the activity of a lanthanide-dependent methanol dehydrogenase - a kinetic and DFT study. Dalton Trans 2018; 47: 10463–10472.

50. Singer H, Steudtner R, Klein AS, Rulofs C, Zeymer C, Drobot B, et al. Minor Actinides Can Replace Essential Lanthanides in Bacterial Life. Angew Chem Int Ed Engl 2023; 62: e202303669.

51. Shin D, Kim J, Kim B-S, Jeong J, Lee J-C. Use of Phosphate Solubilizing Bacteria to Leach Rare Earth Elements from Monazite-Bearing Ore. Minerals 2015; 5: 189–202.

52. Corbett MK, Eksteen JJ, Niu X-Z, Croue J-P, Watkin ELJ. Interactions of phosphate solubilising microorganisms with natural rare-earth phosphate minerals: a study utilizing Western Australian monazite. Bioprocess Biosyst Eng 2017; 40: 929–942.

53. Fathollahzadeh H, Eksteen JJ, Kaksonen AH, Watkin ELJ. Role of microorganisms in bioleaching of rare earth elements from primary and secondary resources. Appl Microbiol Biotechnol 2019; 103: 1043–1057.

54. Prodius D, Klocke M, Smetana V, Alammar T, Perez Garcia M, Windus TL, et al. Rationally designed rare earth separation by selective oxalate solubilization. Chem Commun (Camb) 2020; 56: 11386–11389.

55. Meng X, Zhao H, Zhao Y, Shen L, Gu G, Qiu G. Effective recovery of rare earth from (bio)leaching solution through precipitation of rare earth-citrate complex. Water Res 2023; 233: 119752.

56. Zenker S, Lohmann J, Chiorescu I, Krüger S, Kumke MU, Reich T, et al. Complexation of Ln(III) ions by gluconate: Joint investigation applying TRLFS, CE-ICP-MS, NMR, and DF calculations. Inorg Chem 2025; 64: 7970–7987.

57. Tajmir-Riahi HA. Carbohydrate metal ion complexes. Interaction of D-glucono-1,5-lactone with Zn(II), Cd(II), and Hg(II) ions in the solid and aqueous solution, studied by 13C-NMR, FT-IR, and X-ray powder diffraction measurements. Can J Chem 1989; 67: 651–654.

58. Lumpe H, Daumann LJ. Studies of redox cofactor pyrroloquinoline quinone and its interaction with lanthanides(III) and calcium(II). Inorg Chem 2019; 58: 8432–8441.

59. Lumpe H, Menke A, Haisch C, Mayer P, Kabelitz A, Yusenko KV, et al. The Earlier the Better: Structural Analysis and Separation of Lanthanides with Pyrroloquinoline Quinone. Chemistry 2020; 26: 10133–10139.

60. Gutenthaler-Tietze SM, Mertens M, Phi MT, Weis P, Drobot B, Köhrer A, et al. Comparative binding studies of the chelators methylolanthanin and rhodopetrobactin B to lanthanides and ferric iron. Chembiochem 2026; 27: e202500312.

61. Juma PO, Fujitani Y, Alessa O, Oyama T, Yurimoto H, Sakai Y, et al. Siderophore for Lanthanide and Iron Uptake for Methylotrophy and Plant Growth Promotion in *Methylobacterium aquaticum* Strain 22A. Front Microbiol 2022; 13: 921635.

62. Hofmann M, Retamal-Morales G, Tischler D. Metal binding ability of microbial natural metal chelators and potential applications. Nat Prod Rep 2020; 37: 1262–1283.

63. Pol A, Barends TRM, Dietl A, Khadem AF, Eygensteyn J, Jetten MSM, et al. Rare earth metals are essential for methanotrophic life in volcanic mudpots. Environ Microbiol 2014; 16: 255–264.

64. Cao J, Woodhall MR, Alvarez J, Cartron ML, Andrews SC. EfeUOB (YcdNOB) is a tripartite, acid-induced and CpxAR-regulated, low-pH Fe^2+^ transporter that is cryptic in *Escherichia coli* K-12 but functional in *E. coli* O157:H7. Mol Microbiol 2007; 65: 857–875.

65. Miethke M, Monteferrante CG, Marahiel MA, van Dijl JM. The *Bacillus subtilis* EfeUOB transporter is essential for high-affinity acquisition of ferrous and ferric iron. Biochim Biophys Acta 2013; 1833: 2267–2278.

66. Nakatsuji S, Okumura K, Takase R, Watanabe D, Mikami B, Hashimoto W. Crystal structures of EfeB and EfeO in a bacterial siderophore-independent iron transport system. Biochem Biophys Res Commun 2022; 594: 124–130.

67. Wehrmann M, Berthelot C, Billard P, Klebensberger J. Rare Earth Element (REE)-Dependent Growth of *Pseudomonas putida* KT2440 Relies on the ABC-Transporter PedA1A2BC and Is Influenced by Iron Availability. Front Microbiol 2019; 10: 2494.

68. Chen Z, Lewis KA, Shultzaberger RK, Lyakhov IG, Zheng M, Doan B, et al. Discovery of Fur binding site clusters in Escherichia coli by information theory models. Nucleic Acids Res 2007; 35: 6762–6777.

69. Young GM, Postle K. Repression of *tonB* transcription during anaerobic growth requires Fur binding at the promoter and a second factor binding upstream. Mol Microbiol 1994; 11: 943–954.

70. Reusch RN, Sadoff HL. Putative structure and functions of a poly-beta-hydroxybutyrate/calcium polyphosphate channel in bacterial plasma membranes. Proc Natl Acad Sci U S A 1988; 85: 4176–4180.

71. Reusch RN. Biological complexes of poly-beta-hydroxybutyrate. FEMS Microbiol Rev 1992; 9: 119–129.

72. Reusch RN, Huang R, Bramble LL. Poly-3-hydroxybutyrate/polyphosphate complexes form voltage-activated Ca2+ channels in the plasma membranes of *Escherichia coli*. Biophys J 1995; 69: 754–766.

73. Reusch RN. Transmembrane ion transport by polyphosphate/poly-(R)-3-hydroxybutyrate complexes. Biochemistry 2000; 65: 280–295.

74. Phi MT, Singer H, Zäh F, Haisch C, Schneider S, Op den Camp HJM, et al. Assessing lanthanide-dependent methanol dehydrogenase activity: The assay matters. Chembiochem 2024; 25: e202300811.

75. Ciubotaru I, Biener LC, Vetsova VA, Weis P, Seitz M, Daumann LJ. The elusive chemistry of pyrroloquinoline quinone dimethyl ester lanthanide complexes in biomimetic alcohol oxidation. Eur J Inorg Chem 2025; 28: e202500102.

76. Gu W, Haque MFU, DiSpirito AA, Semrau JD. Uptake and effect of rare earth elements on gene expression in *Methylosinus trichosporium* OB3b. FEMS Microbiol Lett 2016; 363: 1–6.

77. Masuda S, Suzuki Y, Fujitani Y, Mitsui R, Nakagawa T, Shintani M, et al. Lanthanide-dependent regulation of methylotrophy in methylobacteriumaquaticum strain 22A. mSphere 2018; 3.

78. Good NM, Moore RS, Suriano CJ, Martinez-Gomez NC. Contrasting in vitro and in vivo methanol oxidation activities of lanthanide-dependent alcohol dehydrogenases XoxF1 and ExaF from *Methylobacterium extorquens* AM1. Sci Rep 2019; 9: 4248.

79. Chistoserdova L, Kalyuzhnaya MG. Current Trends in Methylotrophy. Trends Microbiol 2018; 26: 703–714.

80. Butterfield CN, Li Z, Andeer PF, Spaulding S, Thomas BC, Singh A, et al. Proteogenomic analyses indicate bacterial methylotrophy and archaeal heterotrophy are prevalent below the grass root zone. PeerJ 2016; 4: e2687.

81. Wilson MC, Mori T, Rückert C, Uria AR, Helf MJ, Takada K, et al. An environmental bacterial taxon with a large and distinct metabolic repertoire. Nature 2014; 506: 58–62.

82. Foerster H-J. The chemical composition of REE-Y-Th-U-rich accessory minerals in peraluminous granites of the Erzgebirge-Fichtelgebirge region, Germany; Part I, The monazite-(Ce)-brabantite solid solution series. Am Mineral 1998; 83: 259–272.

83. Pasero M, Kampf AR, Ferraris C, Pekov IV, Rakovan J, White TJ. Nomenclature of the apatite supergroup minerals. Eur J Mineral 2010; 22: 163–179.

84. Mtyewexr R, Nerer I, Nrcasurue K. A refinement of the crystal structure of gadolinite.

85. Williams CT, Kogarko LN, Woolley AR. Chemical evolution and petrogenetic implications of loparite in the layered, agpaitic Lovozero complex, Kola Peninsula, Russia. Mineral Petrol 2002; 74: 1–24.

86. Shivaramaiah R, Anderko A, Riman RE, Navrotsky A. Thermodynamics of bastnaesite: A major rare earth ore mineral. Am Mineral 2016; 101: 1129–1134.

